# Systematic reduction of the dimensionality of natural scenes allows accurate predictions of retinal ganglion cell spike outputs

**DOI:** 10.1101/2021.10.21.465331

**Authors:** Julian Freedland, Fred Rieke

## Abstract

The mammalian retina engages a broad array of linear and nonlinear circuit mechanisms to convert natural scenes into retinal ganglion cell (RGC) spike outputs. Although many individual integration mechanisms are well understood, predictive models of natural scene encoding perform poorly, likely due to interactions among the active mechanisms. Here, we identified spatial integration mechanisms relevant for natural scene encoding and used computational modeling to predict spike outputs in primate parasol RGCs. Our approach was to simplify the structure of natural scenes and empirically ask whether these changes were reflected in RGC spike responses. We observed that reducing natural movies to 16 linearly integrated regions described ∼80% of the structure of parasol RGC spike responses. We then used simplified stimuli to create high-dimensional metamers that recapitulated the spike response of full-field naturalistic movies. Finally, we identified the retinal computations that convert natural images in 16-dimensional space into 1-dimensional spike outputs.

## Introduction

Visual scenes in the natural world contain complex statistical features that must be reliably encoded by the mammalian visual system to guide behavior (Ruderman and Bialek, 1994). Although physiological mechanisms associated with vision are generally well understood, robust predictions to natural scenes remain elusive (Carandini et al., 2005; Turner et al., 2019). This discrepancy is often attributed to the highly-correlated, non-Gaussian structure of natural scenes that stimulate multiple interacting mechanisms in visual neurons in ways that differ from typical laboratory stimuli.

The spatial complexity of natural scenes is reduced by the retina during the encoding process. Photoreceptors outnumber downstream ganglion cells at ratios of nearly 100:1 (Curcio et al., 1990; Watson, 2014) except in the primate fovea. When paired with classical models that predict linear integration across photoreceptors, encoded scenes lose much of the fine spatial detail that underlies their complex structure (Schreyer and Gollisch, 2021). Indeed, prior work shows that responses of some ganglion cells to natural scenes are well-predicted by linear spatial models (Turner and Rieke, 2016).

Purely linear models, however, do not generalize well across all stimuli and cell types, highlighting the need for new models to capture the encoded structure of scenes (Hochstein and Shapley, 1976; Turner and Rieke, 2016). A rich history of work also identifies situations in which linear integration fails, revealing receptive field subunits that provide sensitivity to spatial structure (Enroth-Cugell and Robson, 1966; Hochstein and Shapley, 1976). Correspondingly, models incorporating nonlinear subunits can improve predicted responses to natural scenes (Euler et al., 2014; Shah et al., 2020; Turner and Rieke, 2016). The impact of subunits on ganglion cell responses can be modulated by activity in the receptive field surround (Enroth-Cugell and Freeman, 1987; Turner et al., 2018), providing a specific example of how interacting mechanisms may shape encoding. Together, these circuit features may be sufficiently generalizable to describe visual stimuli encountered in the natural world, but models of this type have not yet been vetted for their performance.

Here, we tested whether functional subunits and center-surround interactions could account for ganglion cell responses to natural inputs based on the similarity of responses elicited by natural and reduced versions of stimuli. We found that much of the structure of the spike response was retained when natural movies were reduced into eight linearly-integrated regions in both the receptive field center and surround. We confirmed the completeness of this dimensional reduction approach by designing novel high-dimensional metamers that share the same reduced dimensional representation but diverge in other higher-order properties; these spatial metamers elicited similar responses to the original naturalistic scenes. Finally, we probed how stimulus components along these sixteen dimensions were combined to control retinal ganglion cell spike output.

## Results

Our goal was to identify low-dimensional spatial representations of natural stimuli that retained the image structure that shapes responses of On- and Off-parasol retinal ganglion cells (RGCs). We found that we could reduce natural images to 16 spatial regions (i.e. representing a point in a 16-dimensional space) while retaining the majority of structure in the RGC spike response. We then probed interactions between these regions to understand how structure in this 16-dimensional space was converted to a cell’s spike output.

### Spatial integration of natural images in the receptive field center

We started by investigating how the spatial structure of naturalistic scenes in the receptive field center of parasol RGCs could be simplified while minimizing the impact on spike responses. The RGC receptive field is classically modeled as the difference between center and surround components, with the center dominating over some regions of space and the surround others (Fig. 1A) (Hubel and Wiesel, 1959). We measured the center-dominated region of each recorded neuron by first locating the receptive field center and then flashing spatially uniform circular disks of several sizes centered on this location. Responses initially increased with increasing disk size and then decreased as the disks began to impinge on surround-dominated regions. The disk that produced the largest response defined the bounds of the center-dominated region of the neuron’s receptive field.

**Figure 1:**
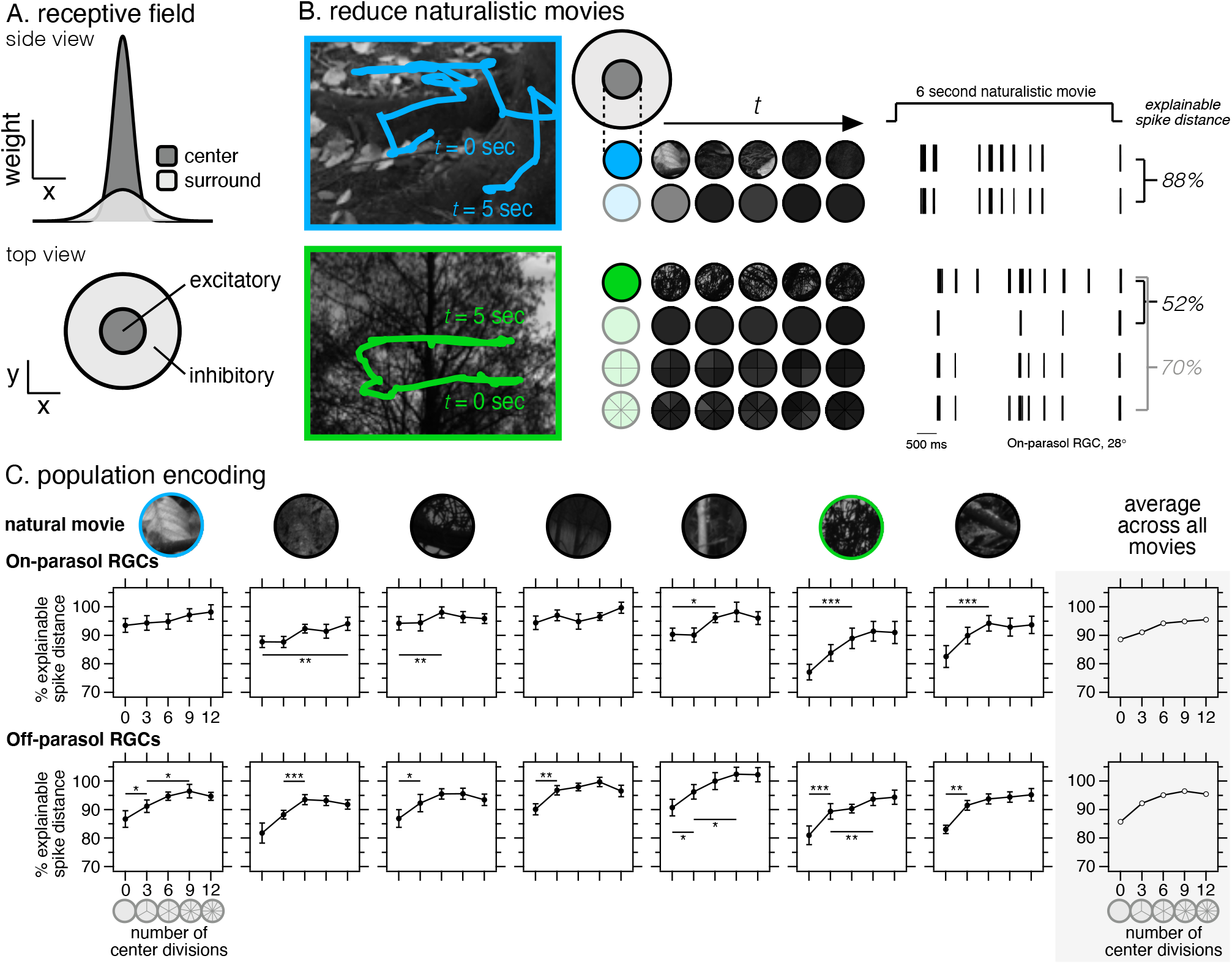
(A) Side- and top-view of a classical 3-D neuronal receptive field. (B, left) Eye-tracking observers during naturalistic image viewing, courtesy of (Van Der Linde et al., 2009), enable the creation of movies that mimic spatiotemporal visual encoding. (B, middle) Each movie frame can subsequently be reduced to a series of linear-equivalent regions. (B, right) Spike traces for a single On-parasol retinal ganglion cell (“RGC”, 28° temporal) in response to various 6-second naturalistic and reduced movies as presented to their receptive field center. Encoding similarities between reduced movies and their naturalistic counterparts are quantified via explainable spike distance (see Methods; (Victor, 2005)). (C) Encoding properties in the receptive field center of individual On- and Off-parasol RGCs (n ≥ 12) as reduced movies are divided into increasing numbers of linear-equivalent regions. Seven unique naturalistic movies, each derived from a unique image, are shown with (right) an average across all moves. Error bars are SEM.

To produce naturalistic stimuli, we used an online database of eye movement trajectories of human observers viewing naturalistic images (Van Der Linde et al., 2009). We used raw recordings of physiological eye trajectories to recreate the scenes that fall on a single neuron during natural viewing. These stimuli (“naturalistic movies”) spanned six seconds in length and were delivered with a circular aperture that restricted them to the receptive field center of the neuron. Each circular naturalistic movie frame was then divided into discrete wedge-shaped regions with each wedge containing a uniform luminance determined by linearly integrating the pixel values of the original movie frame weighted by the estimated receptive field (Fig. 1B; see Equation 1). We quantified the similarity of responses to the original and reduced movies using the explained spike distance (see Methods; (Victor, 2005)). The spike distance measures the similarity of two spike trains by computing a cost to convert one to another. We bounded this distance from above using random spike trains, and from below using the distance between responses to repeated trials of the same natural stimulus. Figure 1B illustrates this process for an example On-parasol RGC. For the top image, the response to the original movie is well approximated by a single spatially uniform region (88% explained spike distance). For the bottom image, the response to the reduced image becomes more similar to that of the original as more regions are added, and correspondingly the explained spike distance increases.

Figure 1C collects the results from this approach across populations of recorded On- and Off-parasol RGCs for seven different images. For Off-parasol RGCs, the explained spike distance consistently increased as the receptive field was subdivided into 6 to 9 regions. This improvement was smaller and more variable for On-parasol RGCs, which is apparent both in the individual images and the average across images (far right panels in Fig. 1C). Previous work found that On-parasol responses to similar natural movies were well approximated by a linear receptive field (Turner and Rieke, 2016); our results here indicate that this spatial linearity holds for some images, while others (e.g. the 6th from the left in Fig. 1C) evoke clear nonlinear spatial integration.

The results illustrated in Figure 1 show that natural movies restricted to the receptive field center of On- and Off-parasol RGCs can be reduced to a 6-9 dimensional space with minimal loss of structure relevant for the cells’ spike responses. We additionally tested movies with circular divisions that divided the receptive field center into a near- and far-center (Fig. S-1, left). We observed that reduced stimuli with eight wedge-shaped regions across the receptive field center performed similarly to stimuli with eight wedge-shaped divisions in both the near- and far-center (for a total of 16 regions). Our observed number of required regions is considerably less than the ∼200 cone photoreceptors and the ∼30 cone bipolar cells that fall within the parasol receptive field (see Discussion).

### Spatial regions combine near-linearly to control RGC response

We next investigated how activity in discrete regions within the receptive field center are integrated to produce the parasol spike output. The results described below support a model in which receptive field subregions are first rectified and then summed to predict the RGC response.

To probe nonlinear behavior, we flashed 48 naturalistic images for 250 ms in the receptive field center of On- and Off-parasol RGCs and measured their spike response (Fig. 2A). As expected, the relationship between spike output and the integrated luminance within the receptive field center was nonlinear for both cell types (Fig. 2B, top). We then computed an effective contrast by first reducing each natural image to 8 subregions, rectifying the values in each subregion, and summing. Spike responses showed a linear or near-linear relationship with this effective contrast (Fig. 2B, bottom), indicating that the effective contrast is a good predictor of RGC spike output.

**Figure 2:**
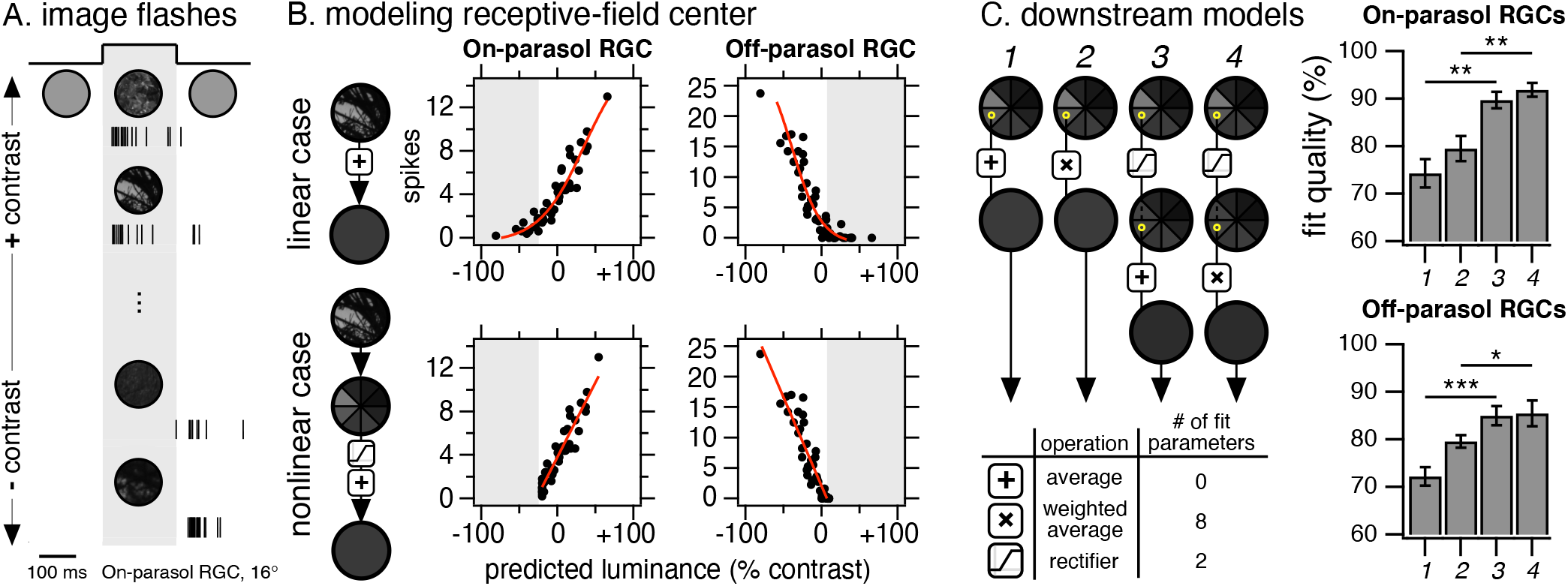
(A) 48 naturalistic images, spanning an array of luminances, were flashed for 250 ms onto the receptive field center of an On-parasol RGC. (B) All 48 naturalistic images were individually integrated through linear and nonlinear models and correlated with spike response. Each point represents the total number of spikes in response to a single naturalistic image flash for one On- and Off-parasol (47°, 16° respectively). (C) Four downstream integration architecture models were fitted to predict spikes responses to 48 naturalistic image flashes in the receptive field center of individual On- and Off-parasol RGCs (n = 9, n = 10, respectively). Percent fit quality was determined by optimizing fit parameters for the best linear correlation (r^2^) between predicted luminance and measured spikes.

To further test this conclusion, we compared the quality of predictions made by four models (Fig. 2C, left): (1) linear summation; (2) weighted linear summation, in which optimal weights were associated with each of the 8 receptive field regions; (3) rectified summation, in which signals in each subregion were rectified and then summed; (4) weighted and rectified summation, in which subregion signals were rectified and then optimally weighted. Figure 2C, right shows the explained variance of the responses across 48 flashed images for each model. Models incorporating rectification performed considerably better than those that did not, and optimizing weights across subregions offered little additional improvement despite the increase in the number of model parameters.

The results illustrated in Figures 1 and 2 are broadly consistent with past work identifying bipolar cells as the origin of receptive field subunits (Demb et al., 1999), with little or no nonlinear interaction between subunits. The size of the subunits, however, is considerably larger than expected for bipolar cells, an observation that we return to in the Discussion.

### Receptive field surrounds have modest impact on responses to naturalistic inputs

We next quantified the impact of the receptive field surround on naturalistic image encoding. To do this, we compared parasol RGC spike responses to naturalistic movies either restricted to the receptive field center or spanning the entire receptive field (Fig. 3A). We used the explained spike distance to quantify differences between responses elicited by center-only and full-field stimulation. Across seven naturalistic movies, receptive field center stimulation alone captured ∼70% of the response to full-field stimuli for On- and Off-parasol RGCs (Fig. 3B). This provides a bound on the impact of the receptive field surround.

**Figure 3:**
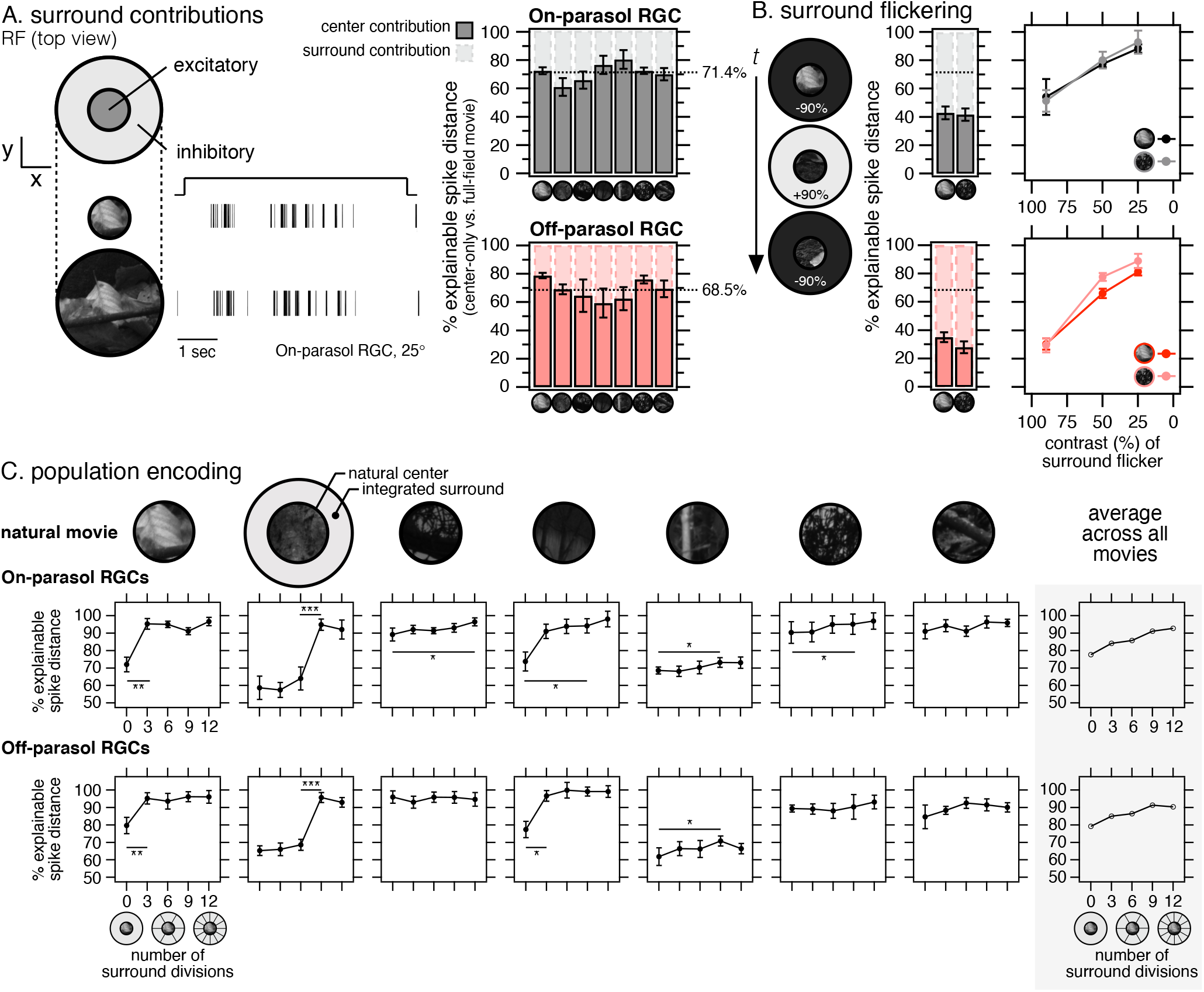
(A) Spike responses to naturalistic movies presented to the receptive field center and full receptive field of an On-parasol RGC (25° temporal). (A, right) Encoding properties of individual On- and Off-parasol RGCs (n ≥ 8) for center-only movies relative to their full-field counterparts. (B, left) Across the same set of On- and Off-parasol RGCs, we presented an artificial movie with a naturalistic center and an artificial, uniform receptive field surround that inverts polarity after every saccade at +/-90% contrast. (B, right) Encoding effects as the contrast of the uniform surround was altered for individual On- and Off-parasol RGCs (n = 4). (C) Encoding properties of individual On- and Off-parasol RGCs (n = 8) to movies with naturalistic centers and reduced linear-equivalent surrounds divided into increasing numbers of regions.

To measure the potential impact of the surround, we measured responses in the same RGCs to movies with a naturalistic image in the receptive field center and an artificial uniform disk in the receptive field surround (Fig. 3B). To stimulate the receptive field surround, the luminance of the surround disk changed polarity after each saccade. We observed that the circuitry of the receptive field surround is capable of imposing a far greater effect on encoding, nearly doubling its impact during full-field encoding (≥ 57%) at 90% contrast (Fig. 3B). The effect of receptive field surround became less pronounced as we decreased the contrast of the surround disk (Fig. 3B). This indicates that the modest impact of the surround during naturalistic movies occurs because such movies do not exercise the surround strongly.

### Spatial integration of natural images in the receptive field surround

Figure 1 shows that the spatial structure in the receptive field center that is relevant to parasol responses is largely captured by dividing the center into eight subregions. The experiments described below indicate that a similar dimensional reduction is possible in the receptive field surround.

We compared spike responses to full-field movies with those to movies with a naturalistic image in the receptive field center and reduced-dimensionality surrounds. The sensitivity of encoding to the number of surround regions varied considerably across images (Fig. 3C); in some cases, dividing the surround into regions had little effect (e.g. rightmost panels), while in others 6 to 9 regions improved the ability to recapitulate responses to the full natural movie. Averaged across images, dividing the surround into regions modestly improved full-field encoding in both On- and Off-parasol RGCs (Fig. 3C, right). Additional experiments with reduced stimuli revealed that eight divisions across the entire receptive field surround performed similarly to stimuli with eight divisions in both the near- and far-surround (totaling 16 regions over the same region of space) (Fig. S-1, right).

### Building low-dimensional full-field reductions

The work described above suggests that approximately eight spatially-uniform regions in both the receptive field center and surround are sufficient to describe parasol responses to a collection of naturalistic stimuli. Together, these studies indicate that any nonlinear aspects of spatial integration occur in a ∼16-dimensional space. Indeed, our 16 reduced-region movies consistently performed better than movies with a single receptive field center and surround (Fig. 4A). Across two naturalistic movies, our 16-D model captured ∼80% of full-field naturalistic spiking in both On- and Off-parasol RGCs (Fig. 4A).

**Figure 4:**
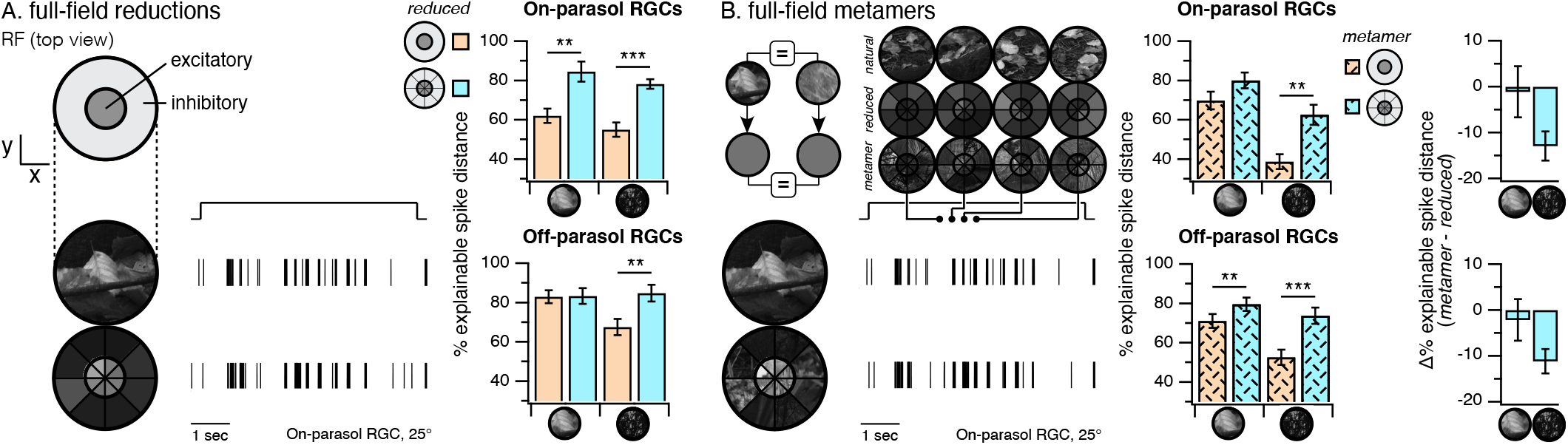
(A) Spike responses to a naturalistic movie and its 16-D reduction as presented to the full receptive field of an On-parasol RGC (25° temporal). (A, right) Encoding properties of individual On- and Off-parasol RGCs (n = 16) for full-field 2-D and 16-D reductions. (B, left) For each reduced linear-equivalent region, we identified sets of naturalistic images that share the same local integrated luminance. Each reduced region was then replaced with its analogous image to produce a metamer. Sample movie frames are included for naturalistic, reduced, and metamer movies at *t* = 2.0, 2.3, 2.7, and 3.0 seconds. 2-D and 16-D metamer movies were then presented to On- and Off-parasol RGCs (n ≥ 15) and compared directly with the original full-field naturalistic movie. (B, right) Change in performance between 16-D metamers and 16-D reductions as averaged across individual cells (n ≥ 15).

### Building high-dimensional full-field metamers

The reduction of high-dimensional stimuli into 16 linear-equivalent regions predicts that different images that share the same 16-dimensional representation should elicit the same response. To test this prediction, we reduced a naturalistic full-field movie to 16-dimensional space and then identified, separately for each of the 16 regions, another image that shared the same integrated luminance. Each reduced region was replaced with its analogous image. Repeating this procedure for each spatial region produces a high-dimensional metamer, consisting of 16 stitched-together image patches from different images that reduce to the same point in 16-dimensional space as the original image (Fig. 4B). This approach was repeated for each full-field naturalistic movie frame. Movie 1 shows a naturalistic movie with its 16-D reduction and 16-D metamer. Although the original and metameric images share a reduced dimensional representation, they diverge across other properties. Hence, a comparison of the responses that they elicit is a strong test of whether the 16-dimension description of the image captures the relevant image features.

Responses to full-field 16-D metamers captured features of the original naturalistic movie nearly as well as the 16-D reduced movies for both On- and Off-parasol RGCs (Fig 4B). The ability of metamers to reproduce responses to the original image declined when fewer than 16 dimensions were included in the reduced space, consistent with the decline in performance of the reduced stimulus itself (Fig 4B; additional data not shown). These results indicate that the additional structure added in constructing the metameric movies has little impact on a cell’s response, confirming that 16-D reductions capture most of the relevant stimulus structure.

### Modeling full-field spatial integration

We next probed how a response in the 16-dimensional space identified above was converted to a spike output. In other words, what is the nonlinear function that operates in the 16-D space? To do this, we extended our center-rectified model (Fig. 2C, model *3*) to include the receptive field surround. We then flashed >60 full-field naturalistic images and recorded the resulting spike responses in individual On- and Off-parasol RGCs.

We investigated a wide variety of surround-focused architectures (Fig. S-2) and evaluated model performance as the correlation (r^2^) of the optimal linear fit between spatially-integrated luminance (output of model) and spike responses across flashed images. The model with the best performance across both On- and Off-parasol RGCs is presented in Fig. 5A. Here, each region in the surround is combined with its nearest center region prior to rectification and integration; this architecture is similar to previously reported center-surround mechanisms (Enroth-Cugell and Freeman, 1987; Turner et al., 2018). The performance of our optimal model is shown for three On- and three Off-parasol RGCs and highlights the predictive capabilities of computationally reducing scenes into 16-D space (Fig. 5B; n > 7). Table 1 contains average nonlinear parameters, as fitted across populations of On- and Off-parasol retinal ganglion cells.

**Table 1:**
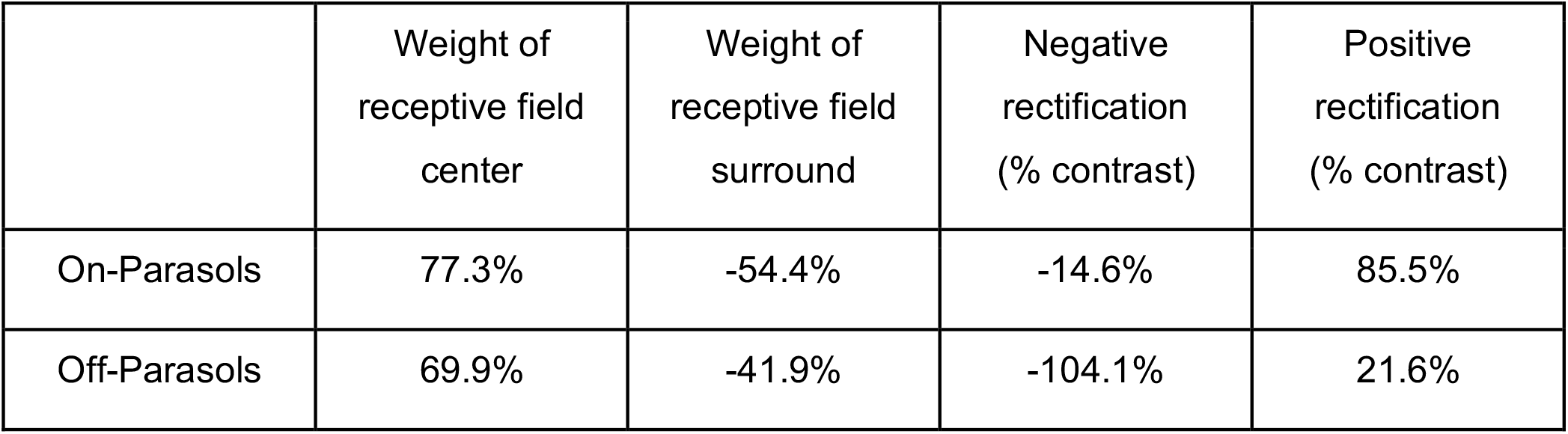
Fit parameters for the full-field modelling architecture presented in Fig. 5A (model *a* in S-2A) as averaged across 9 On- and 7-Off parasol RGCs.

**Table 2:**
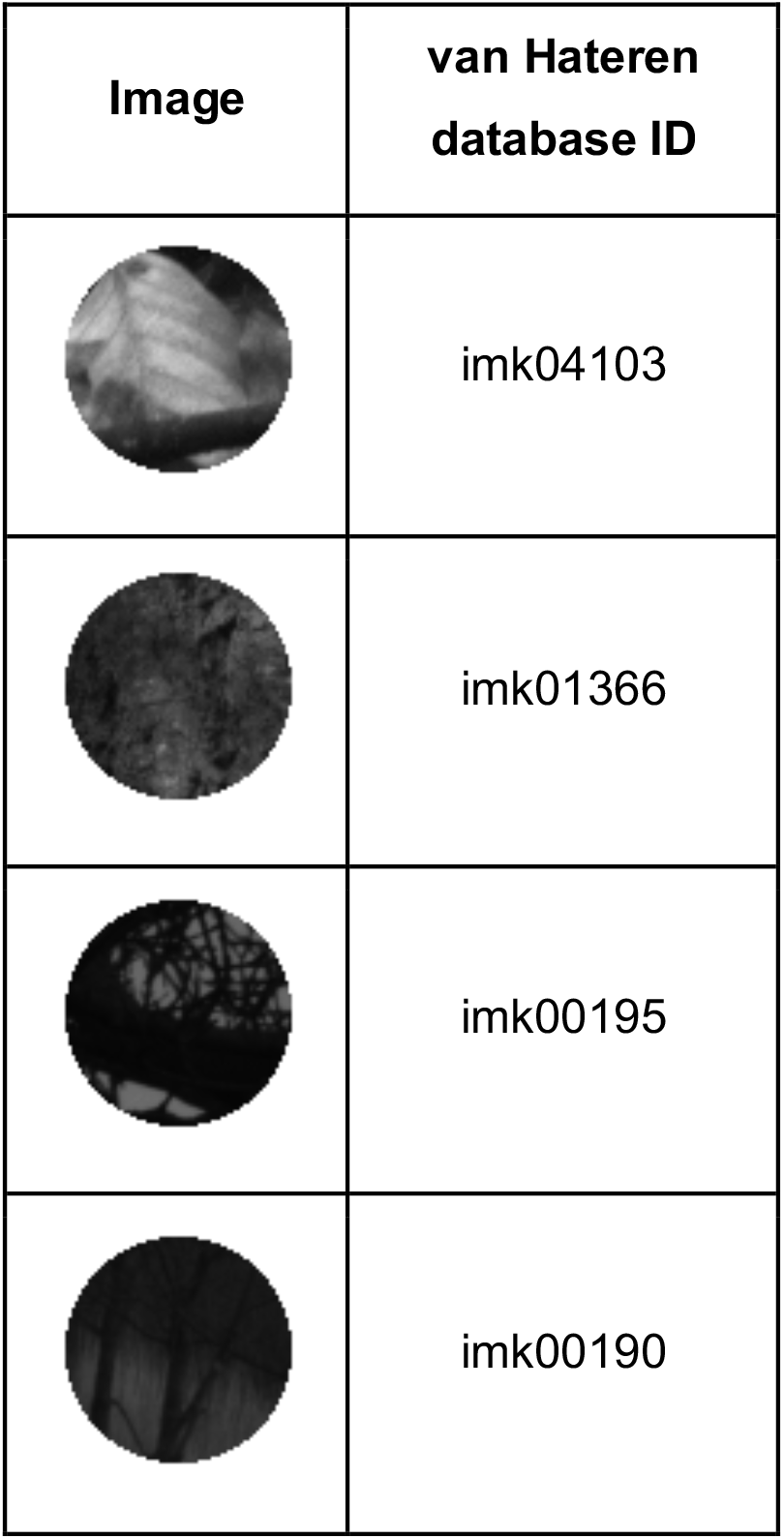
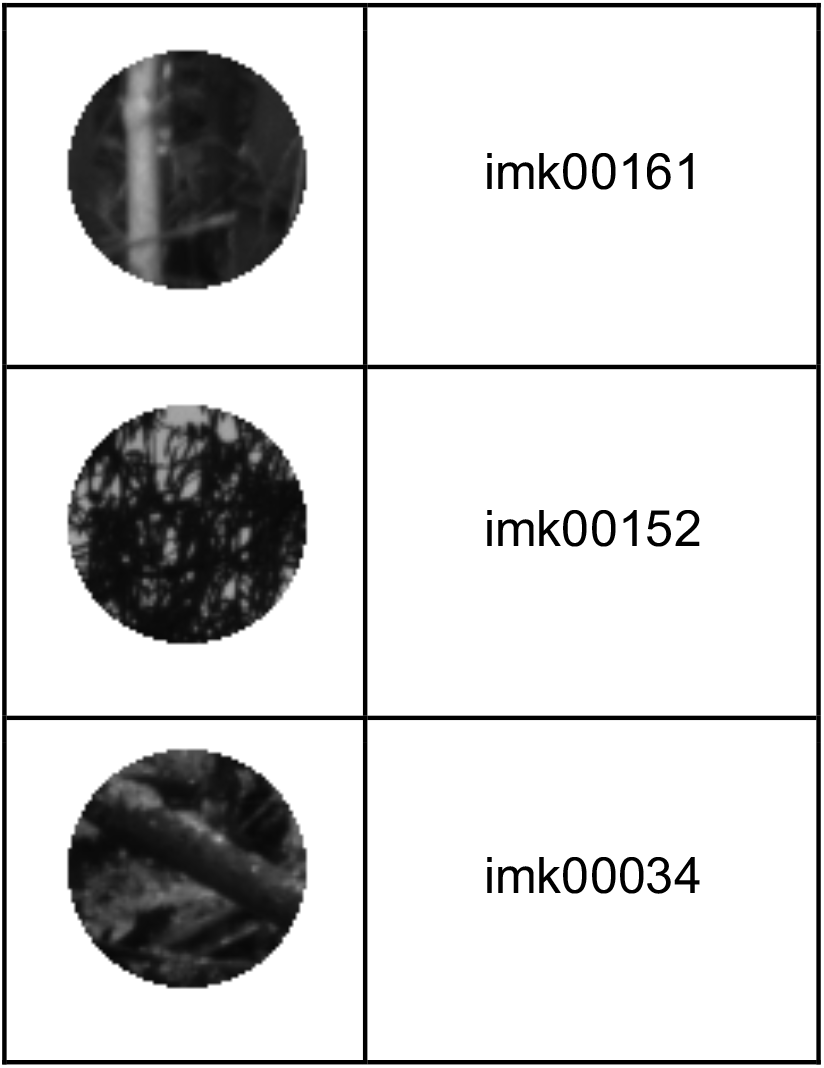
Naturalistic images used in the study and their corresponding ID in the van Hateren database. Top-to-bottom ordering matches left-to-right ordering of images in Fig. 1 and 3.

**Figure 5:**
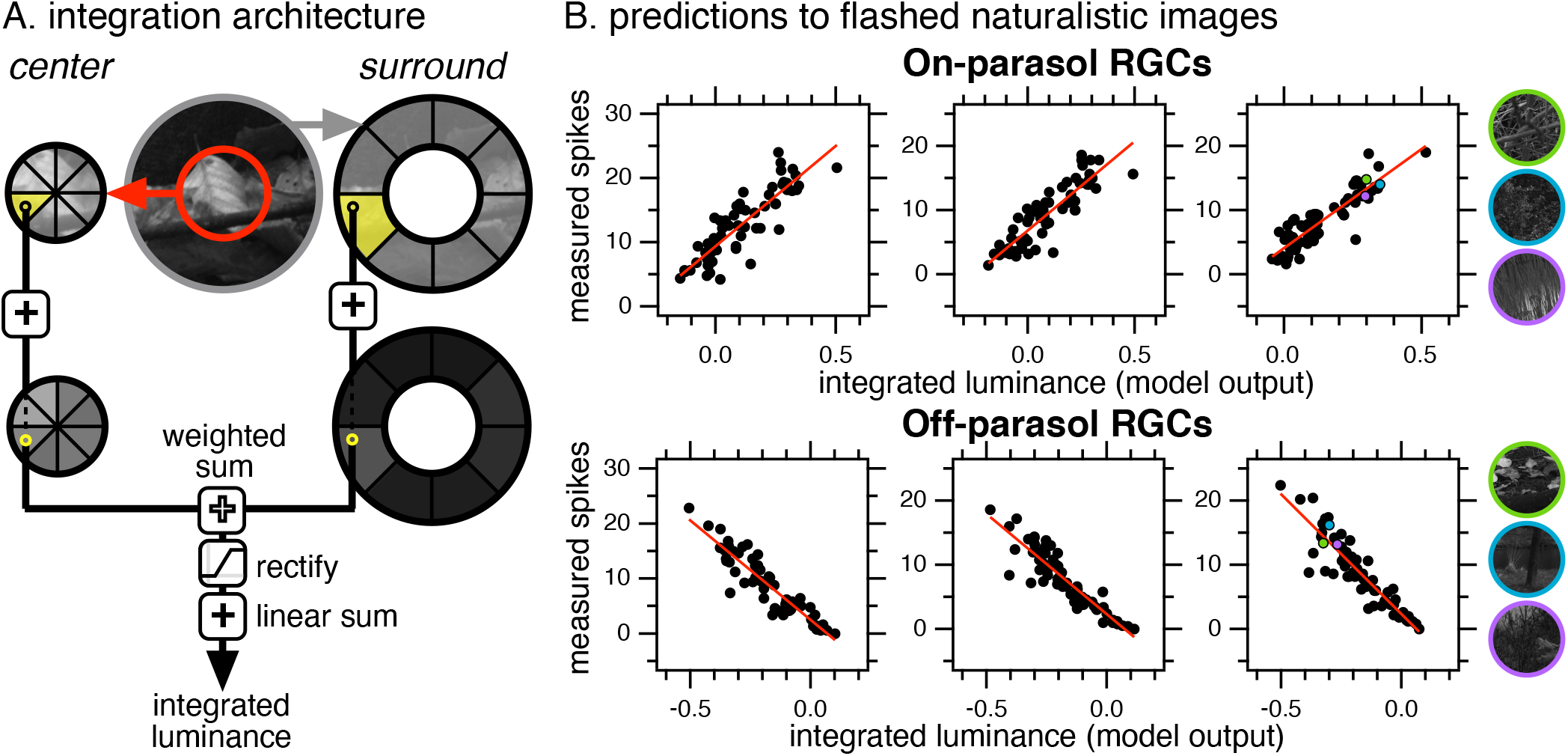
(A) A full-field architecture for uniting spatial integration in the receptive field center and surround for parasol RGCs. The receptive field center and surround are reduced into 8 linear-equivalent regions. The resulting luminances are then uniformly weighted (by one value in the center and another value in the surround). The luminance in each center region is summed with the luminance of the nearest surround region. Resulting values are then rectified and integrated. Other investigated model architectures are available in S-2. (B) Predicted luminances (output of our model) and measured spike responses after flashing 61 - 66 full-field naturalistic images. Three On- and Off-parasol RGCs, selected as the median best-fits in our population of cells, are shown. Three of our ∼60 presented images are highlighted (in green, blue, purple).

The performance of various full-field integration architectures highlighted asymmetries between On- and Off-parasols RGCs (Fig. S-2). On-parasols largely preferred unintegrated surrounds over models with integrated surrounds, with little preference for surround rectification. Off-parasols largely preferred unrectified surrounds over models with rectified surrounds, with little preference for surround integration. Furthermore, On-parasol RGCs preferred the center and surround be integrated before the center was rectified, whereas Off-parasol RGCs did not elicit a clear preference. Our model in Fig. 5A exhibited the best overall performance in On- and Off-parasols for different reasons, adding to literature that suggests significant asymmetries between On and Off pathways (Chichilnisky and Kalmar, 2002; Ratliff et al., 2010; Ravi et al., 2018).

The results in this section indicate that much of the structure of parasol responses to natural image structure can be captured in a two step process: reduce spatial structure by linear integration to a 16-dimensional space, and then combine dimensions in this space in a specific sequence of rectification and integration. This procedure identifies two different types of metamers: pre-integration and post-integration metamers (Fig. 6). Pre-integration metamers capture the luminance encoded by individual subunits and are investigated in Fig. 4B. The metamers reflect the stimulus structure that is discarded by the compression of a spatial image into the 16-dimensional space (see Discussion). Post-integration metamers reflect the mapping of different points in the 16-dimensional subregion space to the same 1-dimensional spike output (Fig. 6B). These two steps together identify the set of images which are predicted to produce the same RGC spike output, and identify how these degeneracies originate.

**Figure 6:**
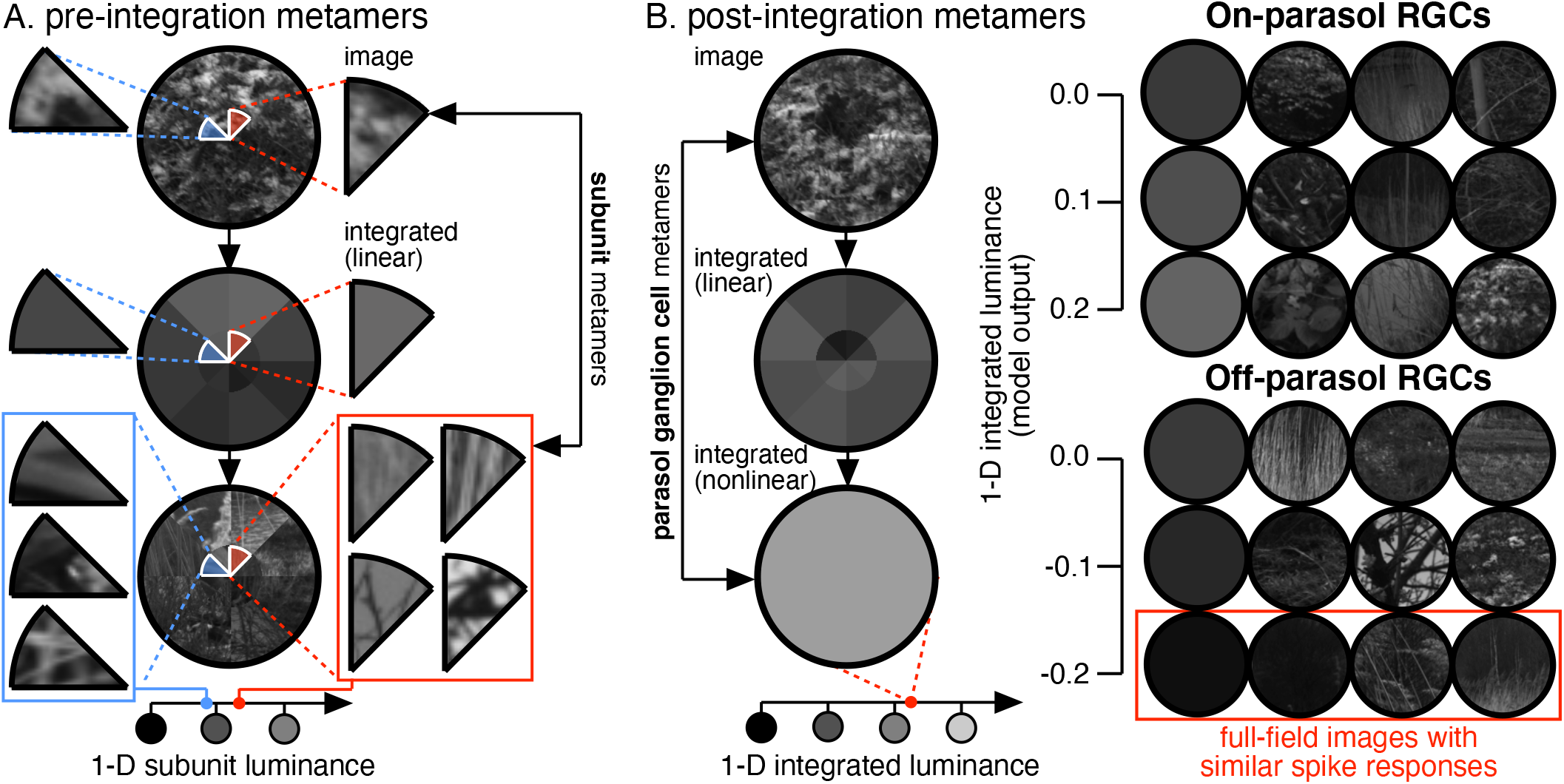
(Left) Pre-integration metamers consist of natural images reduced into 16 linearly integrated regions. Each region is then replaced with a region from another image that reduces to a similar luminance. (Right) Post-integration metamers use fitted nonlinear integration parameters to estimate the 1-dimensional luminance of a natural image. Repeating this operation across a library of natural images reveals sets of full-field images that elicit similar spike responses in On- and Off-parasol RGCs.

## Discussion

We investigated the structure of natural scenes at the level of retinal ganglion cells (RGCs) by comparing the spike response of naturalistic and simplified (“reduced”) naturalistic movies. We observed that reducing movies to eight linearly-integrated spatial regions in the receptive field center and surround captured ∼80% of spiking behavior in On- and Off-parasol RGCs. We then used computational modeling to assess how 16-dimensional movies are integrated into a single 1-dimensional output, and used our best-performing spatial integration model to predict spike responses to flashed full-field natural images.

### On- and Off-parasol retinal ganglion cells reduce the dimensionality of naturalistic movies during spatial integration

Signals from several hundred cone photoreceptors converge through retinal circuits to control the spike output of a single parasol RGC. Understanding how the integration of cone signals works requires identifying linear and nonlinear components of the relevant circuits. Importantly, associativity does not hold across linear and nonlinear mechanisms -- linear integration followed by a nonlinearity is not the same as a nonlinearity followed by linear integration.

Classic center-surround receptive field models assume that linear integration over space and time reduces inputs from a photoreceptor array to a 1-dimensional signal. Linear-nonlinear and generalized-linear models use linear integration as an initial step followed by a single nonlinearity. Decades of physiological studies, however, have identified nonlinear properties of the responses of the photoreceptors and the bipolar cells that convey photoreceptor signals to RGCs (Angueyra and Rieke, 2013; Baylor and Hodgkin, 1973; Demb et al., 2001; Endeman and Kamermans, 2010; Enroth-Cugell and Robson, 1966; Hochstein and Shapley, 1976). Such nonlinear behaviors can increase sensitivity to spatial structure on a scale considerably smaller than predicted by the classical receptive field.

We determined empirically how the division of the receptive field into subregions impacted spike responses; each subregion that we defined integrated inputs linearly. We found that naturalistic inputs could be reduced to sixteen linear subregions with minimal changes in spike response. The performance of metamers (Fig. 4B, see Mov. 1) relative to their naturalistic counterparts provides a direct test of the completeness of this space: the original movie and matched metamer share little other than the same reduced dimensional representation, yet elicit very similar spike responses across a 6-second movie. This highlights that we have identified a reduced dimensional space in which the nonlinear operations that control spike output operate and enabled the testing of different sequences of nonlinear steps that convert a point in this space into a RGC’s 1-dimensional spike output (Fig. 6A).

The dimensional reduction of images presents an opportunity to substantially simplify and study the underlying dimensionality of scenes. Computations that reduce natural images into 16 dimensional space amplify certain spatial scales over others. Additionally, adjacent linear regions remain highly correlated to each other, conserving some of the correlation structure of the original image. This suggests that some of the technical difficulties associated with analyzing natural images are still conserved in our 16-dimensional model. In the context of work that seeks to distinguish between natural and artificial stimuli, correlations within our 16-dimensional space may provide an avenue for analysis.

Rectification in the receptive field center of our reduced model appears to mimic the rectification known to occur at the bipolar cell synapse (Demb et al., 2001). However, since the receptive field center of a parasol RGC typically spans a diameter of 200 to 300 μm, a single bipolar cell would need to span a diameter of 70 to 100 μm to encode the same number of pixels as a single subregion. This agrees with the functional size of diffuse bipolar cells (∼90μm) but diverges from their anatomical size (30 to 50 μm) (Dacey et al., 2000). Thus, our model appears to cluster together multiple bipolar cells in single functional subunits. This is consistent with computational approaches to identify subunits, which similarly find that responses in the receptive field center are well described by as few as five functional subunits (Shah et al., 2020). Why functional subunits appear so much larger than expected from anatomy is unclear.

The shape and distribution of reduced subregions are also of curiosity. Our reduced model bins an array of subunits into discrete spatial wedge-shaped clusters that do not have a physiological basis. This suggests that there is some flexibility in determining subunit shape and location.

### Reduced images reveal spatial integration mechanisms in On- and Off-parasol retinal ganglion cells

We observed that receptive field surrounds had a modest impact on coding of natural images (Fig. 3B). Surrounds in the same cells were strong for artificial stimuli, and hence the modest impact of surrounds for coding of natural images occurred because these images did not activate the surround strongly. This effect was also observed during our flashed full-field image patches. Full-field flashed images that we predicted to only contain rectification in the receptive field center could be modeled with >70% performance using center-only models (Fig. S-2C, “center-only models”). Only images that contained rectification in the surround distinguished strongly between surround models.

The performance of surround models across populations of RGCs highlighted previously reported asymmetries between On and Off pathways (Chichilnisky and Kalmar, 2002; Ratliff et al., 2010; Ravi et al., 2018). Only On-parasols preferred that the receptive field surround remain divided into eight linear regions during center-surround integration. Rectification in the receptive field surround only resulted in a large decrease in performance for Off-parasols. This suggests that our optimal full-field model (Fig. 5A), which yielded the best performance in both On- and Off-parasols, captured different integration mechanisms across pathways.

The best universal full-field model (Fig. 5A) operates in four stages. First, the center and surround are each reduced into eight regions according to receptive field properties of the neuron. Luminances in center and surround are each uniformly weighted. (For example, using averaged values in Table 1, all luminance values in the receptive field center and surround of a On-parasol would be weighted by 77% and -54%, respectively). The weighted value of each surround region is then added to the weighted value of the nearest center region. The eight resulting values are then rectified and linearly averaged to produce one single luminance value. This model requires four fitted parameters: weights in the center and surround and upper and lower bounds for rectification. These parameters can be used to identify sets of full-field metamers that we expect will elicit similar responses (Fig. 6B).

## Methods

### Generating naturalistic stimuli

Spatiotemporal stimuli were generated using eye motion trajectories of freely-moving observers viewing still naturalistic images (Van Der Linde et al., 2009). Each frame was downsampled to 60 Hz and segmented into spatial regions. For each pixel *p* in region *r* of image *I*, the linear-equivalent luminance, *L*, was calculated via,

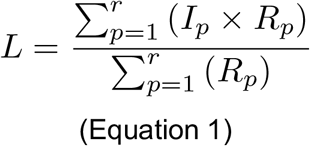

where *R* is the receptive field of an individual neuron.

Metamers were generated by calculating linear-equivalent luminance across a library of 1826 patches sampled from natural images. The patch which had a linear-equivalent luminance closest to that of the original movie was selected. This process was repeated independently for each spatial region. The resulting spatial metamers included a collection of patches from multiple images. A MATLAB package for generating naturalistic movies, reductions, and metamers is available on GitHub: https://github.com/jfreedland/reduce-natural-scenes.

### Explainable spike distance

Spike responses for reduced stimuli were compared to naturalistic responses via explainable spike distance. The spike distance quantifies the cost for adjusting a spike train response for an experimental stimulus (reduced movie) into a spike train response for a control stimulus (naturalistic movie) (Victor, 2005). The two spike trains are matched by a combination of shifting spikes in time, with a cost proportional to the time shift, and deleting and adding spikes. To calculate the explainable spike distance, we compared the calculated spike distance with an upper bound formed from random spike trains with the same spike count, and a lower bound formed from spike responses to repeats of the same stimulus. If the spike distance for a given reduced image equaled the lower bound (corresponding to 100% explainable spike distance), the responses to that stimulus were indistinguishable from those to the original stimulus. If the spike distance equaled the upper bound (corresponding to 0% explainable spike distance), none of the structure of the response to the original stimulus was retained in the response to the reduced stimulus.

### Stimulus projection

Pre-generated .mp4 movies were focused onto RPE-attached macaque photoreceptors via an OLED microdisplay monitor (eMagin). Each monitor pixel (600 × 800 pixels) corresponded to 1.65 μm or 0.5 arcmin. Presented movies (300 × 400 pixels) were enlarged to match the spatial scale of physiological eye movements from the DOVES database (1 pixel per 1 arcmin) (Van Der Linde et al., 2009). The average pixel intensity across naturalistic images used in the study are similar (16.3-18.6% of total monitor intensity); 17% monitor intensity was scaled to correspond to 5000 L/M-cone isomerizations per second. Monitor outputs were linearized via gamma correction. Stimuli presentation and data acquisition was guided by MATLAB software packages Stage (http://stage-vss.github.io) and Symphony (http://symphony-das.github.io).

### Cell identification & acquisition

Retinal ganglion cells were initially categorized by soma size, receptive field size, and spike transience in response to a step in light intensity. Cell type was confirmed by observing unique responses to naturalistic movies. Recorded cells were screened for sensitivity, and data was collected only from cells that exhibited at least 20 sp/s modulation of their firing rate in response to a uniform 5% contrast, 4 Hz-modulated stimulus at 5000 R^*^/cone/s.

## Supplemental Figures / Tables

**Figure S-1:**
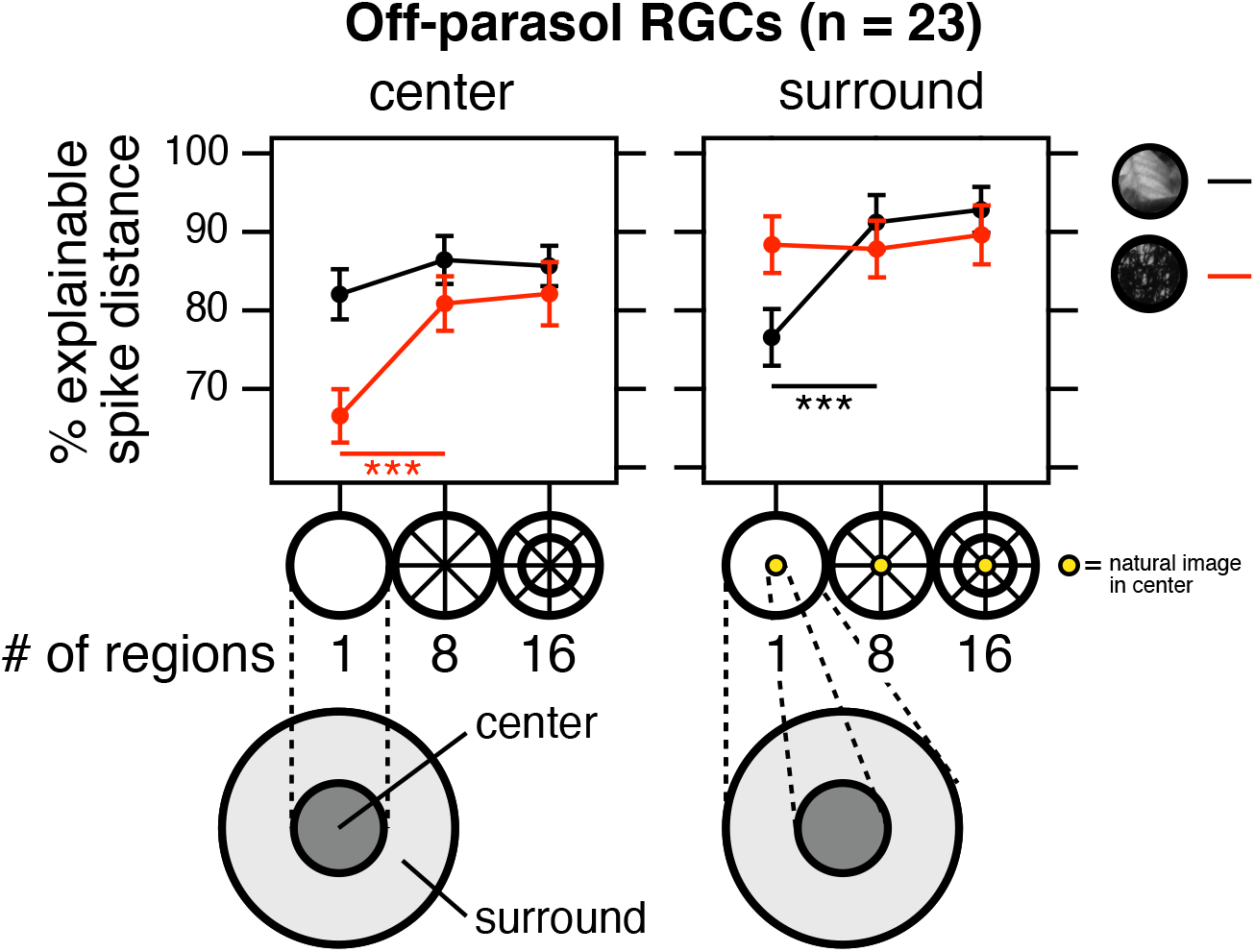
Explainable spike distance for Off-parasol RGCs (n = 23) upon presenting two natural image movies with reduced linear regions in the receptive field (left) center or (right) surround. Reduced movies with 1 and 8 regions are similar to stimuli in Fig. 1C (left) and Fig. 3C (right). Reduced movies with 16 regions contain eight divisions in both the near- and far-center/surround. The near-center/surround is defined as a spatial region that captures 50% of measured excitatory/inhibitory behavior.

**Figure S-2:**
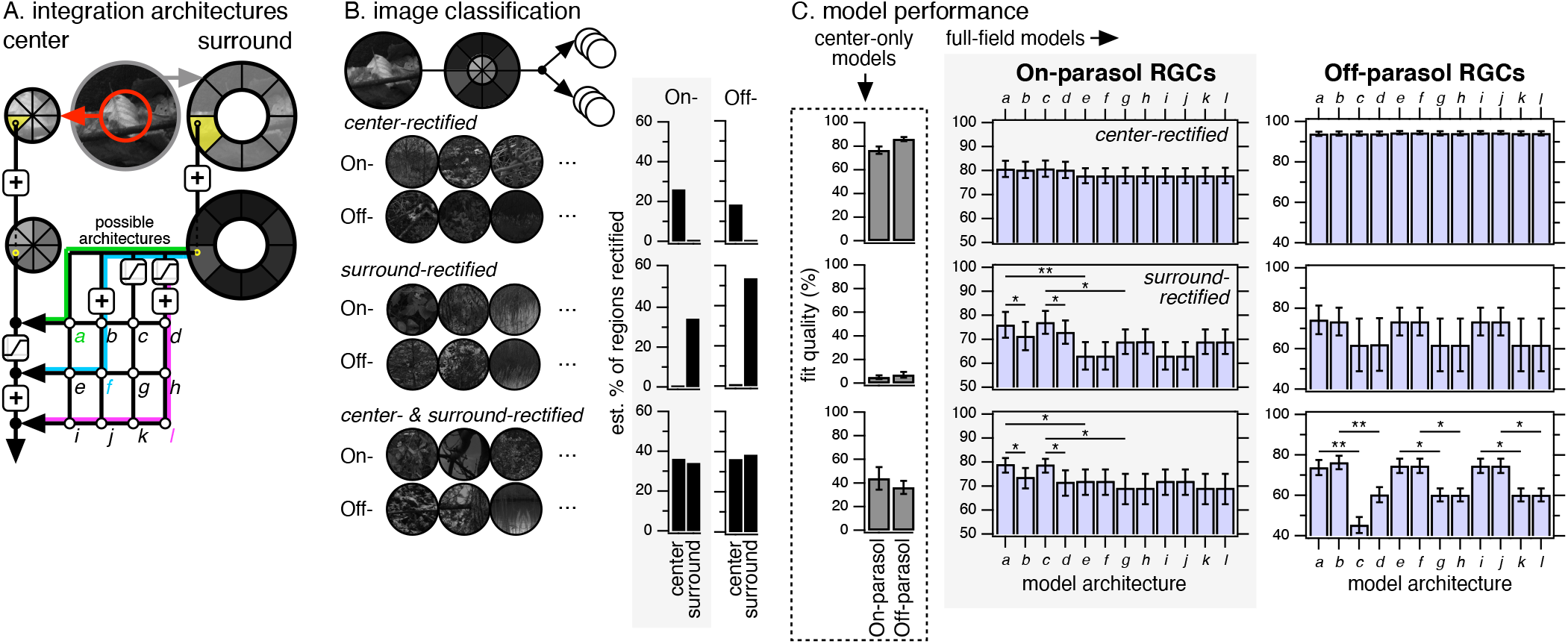
(A) Set of possible architectures for uniting spatial integration in the receptive field center and surround for parasol RGCs. Receptive field surrounds were reduced into 8 linear-equivalent regions and added to the receptive field center pre-rectification, post-rectification, or post-integration after undergoing a set of operations (rectification, integration, neither, or both). This produced 12 total combinations of center-surround architectures; three such models are highlighted (*a, f, l*). Model *a* is equivalent to the model in Fig. 5A. (B) For model testing, we identified three different collections of naturalistic images (16-25 images per dataset) for On- and Off-parasols that share similar amounts of rectification in the receptive field center and/or surround. (C) All three sets of naturalistic images were flashed (for 250 ms) onto individual On- and Off-parasol RGCs (n = 7, n = 9, respectively) and modeled through each downstream circuit architecture. Center-only models were calculated by ignoring surround luminances and fitting our best center model (Model 3 in Fig 3C).

**Movie 1:**
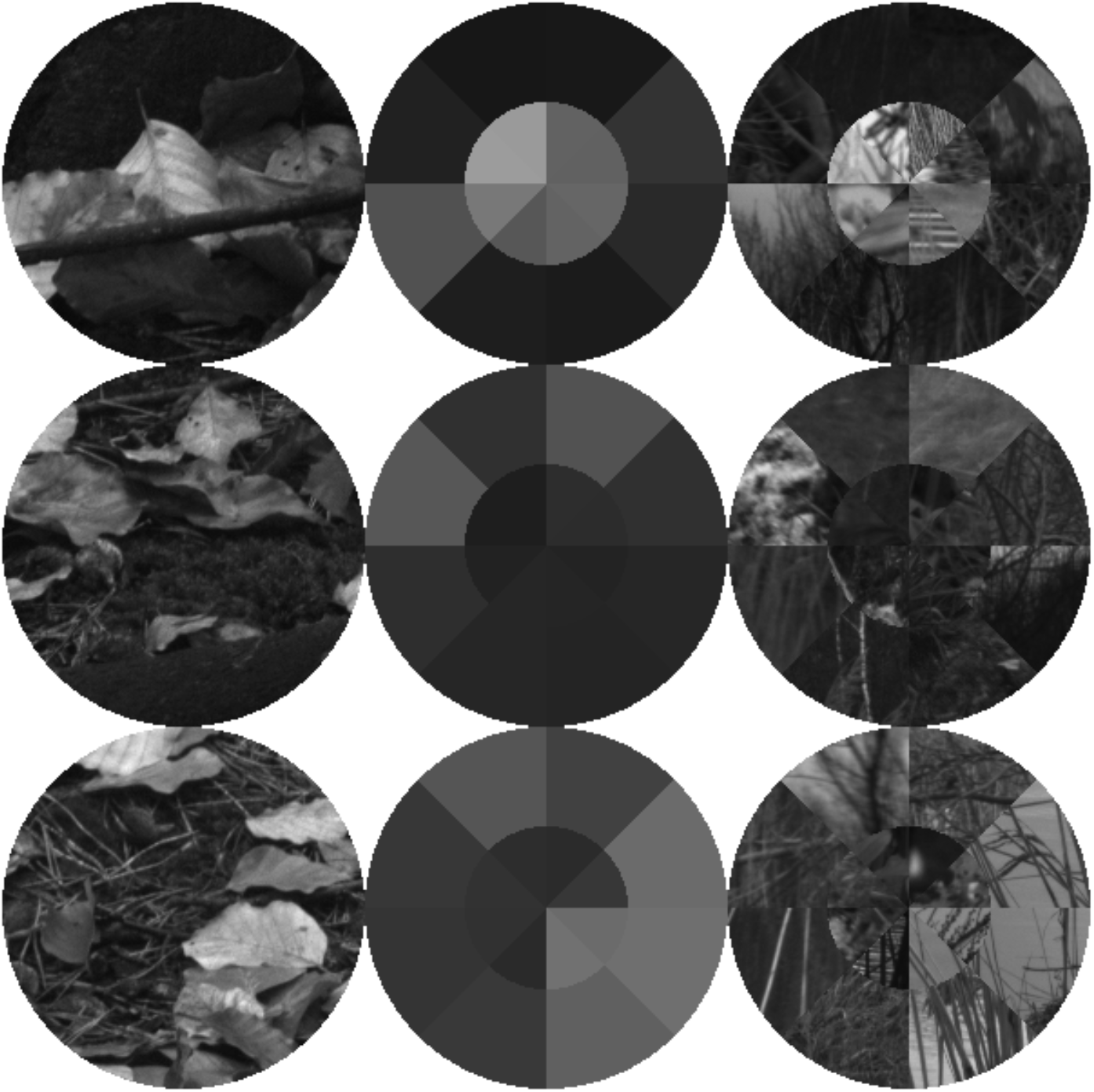
For this preprint, we have isolated three movie frames (from top to bottom: *t* = 1, 2, 3 seconds). (Left) A sample naturalistic movie (“imk04103”) generated using human eye movements. (Middle) imk04103 reduced into full-field 16-D parasol space. (Right) A full-field 16-D metamer that reduces into an equivalent space as imk04103.

## References

Angueyra, J.M., and Rieke, F. (2013). Origin and effect of phototransduction noise in primate cone photoreceptors. Nat. Neurosci. 16, 1692–1700.

Baylor, D.A., and Hodgkin, A.L. (1973). Detection and resolution of visual stimuli by turtle photoreceptors. J. Physiol. 234, 163–198.

Carandini, M., Demb, J.B., Mante, V., Tolhurst, D.J., Dan, Y., Olshausen, B.A., Gallant, J.L., and Rust, N.C. (2005). Do We Know What the Early Visual System Does? J. Neurosci. 25, 10577–10597.

Chichilnisky, E.J., and Kalmar, R.S. (2002). Functional asymmetries in ON and OFF ganglion cells of primate retina. J. Neurosci. Off. J. Soc. Neurosci. 22, 2737–2747.

Curcio, C.A., Sloan, K.R., Kalina, R.E., and Hendrickson, A.E. (1990). Human photoreceptor topography. J. Comp. Neurol. 292, 497–523.

Dacey, D., Packer, O.S., Diller, L., Brainard, D., Peterson, B., and Lee, B. (2000). Center surround receptive field structure of cone bipolar cells in primate retina. Vision Res. 40, 1801–1811.

Demb, J.B., Haarsma, L., Freed, M.A., and Sterling, P. (1999). Functional Circuitry of the Retinal Ganglion Cell’s Nonlinear Receptive Field. J. Neurosci. 19, 9756–9767.

Demb, J.B., Zaghloul, K., Haarsma, L., and Sterling, P. (2001). Bipolar Cells Contribute to Nonlinear Spatial Summation in the Brisk-Transient (Y) Ganglion Cell in Mammalian Retina. J. Neurosci. 21, 7447–7454.

Endeman, D., and Kamermans, M. (2010). Cones perform a non-linear transformation on natural stimuli. J. Physiol. 588, 435–446.

Enroth-Cugell, C., and Freeman, A.W. (1987). The receptive-field spatial structure of cat retinal Y cells. J. Physiol. 384, 49–79.

Enroth-Cugell, C., and Robson, J.G. (1966). The contrast sensitivity of retinal ganglion cells of the cat. J. Physiol. 187, 517–552.

Euler, T., Haverkamp, S., Schubert, T., and Baden, T. (2014). Retinal bipolar cells: elementary building blocks of vision. Nat. Rev. Neurosci. 15, 507–519.

Hochstein, S., and Shapley, R.M. (1976). Quantitative analysis of retinal ganglion cell classifications. J. Physiol. 262, 237–264.

Hubel, D.H., and Wiesel, T.N. (1959). Receptive fields of single neurones in the cat’s striate cortex. J. Physiol. 148, 574–591.

Ratliff, C.P., Borghuis, B.G., Kao, Y.-H., Sterling, P., and Balasubramanian, V. (2010). Retina is structured to process an excess of darkness in natural scenes. Proc. Natl. Acad. Sci. U. S. A. 107, 17368–17373.

Ravi, S., Ahn, D., Greschner, M., Chichilnisky, E.J., and Field, G.D. (2018). Pathway-Specific Asymmetries between ON and OFF Visual Signals. J. Neurosci. 38, 9728–9740.

Ruderman, D.L., and Bialek, W. (1994). Statistics of natural images: Scaling in the woods. Phys. Rev. Lett. 73, 814–817.

Schreyer, H.M., and Gollisch, T. (2021). Nonlinear spatial integration in retinal bipolar cells shapes the encoding of artificial and natural stimuli. Neuron 109, 1692-1706.e8.

Shah, N.P., Brackbill, N., Rhoades, C., Kling, A., Goetz, G., Litke, A.M., Sher, A., Simoncelli, E.P., and Chichilnisky, E. (2020). Inference of nonlinear receptive field subunits with spike-triggered clustering. ELife 9, e45743.

Turner, M.H., and Rieke, F. (2016). Synaptic Rectification Controls Nonlinear Spatial Integration of Natural Visual Inputs. Neuron 90, 1257–1271.

Turner, M.H., Schwartz, G.W., and Rieke, F. (2018). Receptive field center-surround interactions mediate context-dependent spatial contrast encoding in the retina. ELife 7, e38841.

Turner, M.H., Sanchez Giraldo, L.G., Schwartz, O., and Rieke, F. (2019). Stimulus- and goal-oriented frameworks for understanding natural vision. Nat. Neurosci. 22, 15–24.

Van Der Linde, I., Rajashekar, U., Bovik, A.C., and Cormack, L.K. (2009). DOVES: a database of visual eye movements. Spat. Vis. 22, 161–177.

Victor, J.D. (2005). Spike train metrics. Curr. Opin. Neurobiol. 15, 585–592.

Watson, A.B. (2014). A formula for human retinal ganglion cell receptive field density as a function of visual field location. J. Vis. 14, 15.

